# A High Accurate Machine Learning Meta-Strategy for the Prediction of Intrinsically Disorder Proteins

**DOI:** 10.1101/2020.05.18.103200

**Authors:** Chengbin Hu, Yiru Qin, Chuan Ye, Jiao Jin, Ting Zhou, Zhixu He, Liping Shu

## Abstract

**Background:** Many proteins or partial regions of proteins do not have stable and well-defined three-dimensional structures *in vitro*. Understanding intrinsically disorder proteins (IDPs) is significant for interpreting biological function as well as studying many diseases. Although more than 70 disorder predictors have been invented, many existing predictors are limited on the characteristics of proteins and do not have very high accuracy. Therefore, it is critical to formulate new strategies on disorder protein prediction.

**Results:** Here, we propose a machine learning meta-strategy to improve the accuracy of disordered proteins and disordered regions prediction. We first use logistic forward parameter selection to select eight most significant predictors from the current available IDP predictors. Then we design a novel meta-strategy using several machine learning models, including Decision-tree based algorithm, Naive Bayes, Random forest, and Convolutional Neural Network (CNN). By applying different strategies, the results suggest Random forest can improve the predicted single amino acid accuracy significantly to 93.35%. Using the combination vector data of eight most significant predictors as input, the Convolution Neural Network can improve the whole protein prediction to 95.62%.

**Conclusion:** According to the performance of our machine learning meta-strategy, the Random forest and CNN models can improve the accuracy to predict intrinsically disorder proteins.

## Background

Around 70% of Protein Data Bank (PDB) structures have some disordered residues, as well as about 25 percent have intrinsically disorder regions (IDRs) more than 10 residues in length [1]. The system studies of intrinsically disorder proteins (IDPs), not only revealed the abundance of IDRs but also linked IDRs with many diseases, including cancer, cardiovascular diseases, amyloidosis, neurodegenerative diseases, diabetes, etc. [2]. There are many computational methods widely used in the study of IDPs and IDRs because they are more efficient and more economy compared with traditional experimental methods [3]. To date, more than 70 disorder protein predictors have been invented [4]. However, they are challenged by limitation of protein characteristics and the increasing number of protein datasets [5]. One possible strategy to solve this problem is to use meta-strategy based methods to combine current exist individual predictors, because meta-strategy is able to combine those “orthogonal true predictions” and improve the final true prediction rate [6]. In addition, meta-strategies have been used in multiple field, such as protein fold recognition, protein secondary structure prediction, protein interaction, protein subcellular locations, post-translational modification, promoter prediction, and many others [7, 8]. Therefore, we plan to develop a novel meta-strategy by integrating the individual predictors that have top performance, including IUPred, DisEMBL, Espritz, RONN, PONDR@VXLT, PONDR-VSL2, Globplot, and Dispro [9, 10, 11, 12]. The PONDR-VSL2, DisEMBL, and Globplot are based on artificial neural network [9, 10]; PONDR@VLXT is based on support vector machines [9]; Dispro used a combination of neural networks and Bayesian methods [9]; IUPred, Espritz and RONN have been developed by both physics-based methods and neural network [10, 11]. We first selected most popular predictors from those predictors that are widely used in scientific research and have well-maintained software packages or web servers. We then select most significant predictors by performing regression analysis. We integrate the predictors that has the best performance into our meta-strategy predictor. A machine learning based classification model is built for classify and predict disordered amino acid. In addition, we apply the Convolutional Neural Network model to analyze the entire protein data and classify the proteins as several classes of disordered protein.

Many popular machine learning techniques are now used in protein prediction, Lee et al. [13] proposed an alternating decision tree algorithm for assessing protein interaction reliability. Geng et al. [14] proposed the method using Naive Bayes to predict Protein-Protein interaction sites. Qi et al. [15] present a new method by constructing random forest to compute the similarities of protein to protein to classify pairs of proteins as interacting or not. Ramkumar et al. [16] proposed the multilayer perceptron approach to predict protein secondary structures using different set of input features and network parameters in distributed computing environment.

Convolutional Neural Network (CNN) is widely used as image recognition classifier because of its high performance for image data [17]. Recently, scientist also use CNN on protein prediction such as DNA–protein binding[18]. Haoyang Zeng et al. [19] identified the best CNN architectures by varying CNN parameters, depth and pooling designs. Because biological data can be very huge, CNN could be a good solution for classifying these data. In David R. Kelley et al.’s study [20], they trained a compendium of accessible genomic sites with DNase-seq data which yields a high accuracy than previous methods. To use CNN to process protein data, we can consider a protein data sequence as an image. Instead of processing image pixel data, we process a sequence of metrics of protein. We also can arrange the 1-D sequence of protein metrics to 2-D matrix data for CNN’s processing.

In this paper, we test the accuracy of different classifiers for single amino acid. We also rearranged the protein data to a matrix for CNN classification. We identify eight most significant individual disordered protein predictors that can build high accuracy meta-strategy prediction models. Using combination of the eight individual disordered protein predictors as input. We propose a machine learning model that has 93.35% accuracy to classify single amino acid. By deploying a matrix transforming and combination for a whole protein, we designed an input of each protein that can be classified easily for Convolutional Neural Network model. We build a two-layer CNN model that has accuracy 95.62% for the whole protein data classification. Our results shows a significant improvement of the prediction of the intrinsically disorder proteins

## Results

### The selection of individual predictors

There are many published IDPs predictors with aviaiable web servers as we introduction in background. The first step is to find the most significant predictors from the current predictors. After literature review and web server checking, we got 11 predictor for analysis as shown in Table 1. We first test the each individual predictor’s performance by check accuracy and true positive rate. Figure 3 shows the fraction of true positive rate and the fraction of true negative rate. We can find these individual predictors have true positive rate range from 35.3% to 79.6%. We further check the accuracy of a single predictor on our data as shown in Table 1.The accuracy of all of these predictors are around 65.2%-77.3%, which is not high enough to trust. Therefore we perform a logistic forward selection to select the most significant predictors as we described in section 2.2. The step by step sequence is shown as the row sequence in Table 3. IUPred is the most significant predictor for the model and is first selected, followed by Disemble, Espritz and so on. The contribution of each predictor to the model equation 1 can be calculated by the the Model *R*^2^. We found that after the logistic model includes the first 8 predictors, the Model *R*^2^ reached the peak value as 0.8731. If we manually include other 3 predictors into the model, the the Model *R*^2^ will consistently decrease as shown in Table 1, which suggest last three predictors have negative interaction with the eight predictors before. Therefore, we use the first 8 predictors to build high accuracy machine learning models.

**Table 1.**
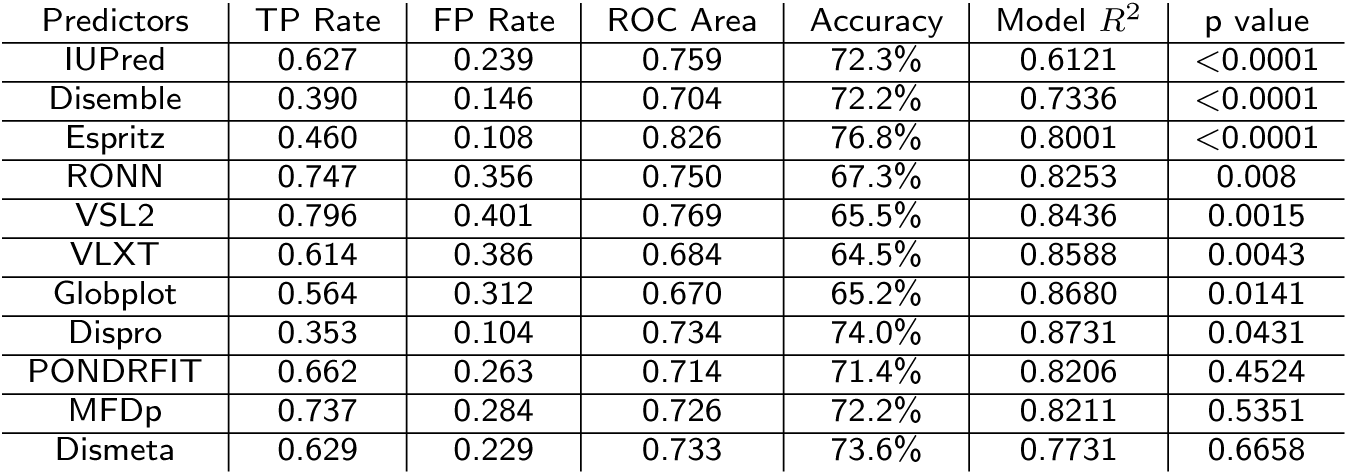
Comparison of current predictors

**Table 2.** Confusion Matrix of single amino acid classifiers

| Classifiers | J48 |  | Naive Bayes |  | Random Forest |  | Multilayer Perceptron |  |
| --- | --- | --- | --- | --- | --- | --- | --- | --- |
| Results | D | O | D | O | D | O | D | O |
| D | 58372 | 33988 | 60111 | 32249 | 71199 | 21161 | 43698 | 48662 |
| O | 16432 | 299899 | 57359 | 258972 | 6018 | 310313 | 24357 | 291974 |

D: Amino acid belongs to Disorder region
O: Amino acid belongs to Ordered region
The second row shows the label for the results of single classifier. The first column shows the ground truth label.

**Table 3.** Comparison of different single amino acid classifiers

| Classifier | TP Rate | FP Rate | Precision | Recall | F-Measure | MCC | ROC Area | Accuracy |
| --- | --- | --- | --- | --- | --- | --- | --- | --- |
| J48 | 0.877 | 0.297 | 0.872 | 0.877 | 0.872 | 0.627 | 0.873 | 87.67% |
| NB | 0.781 | 0.311 | 0.804 | 0.781 | 0.789 | 0.434 | 0.814 | 78.07% |
| RF | 0.933 | 0.182 | 0.933 | 0.933 | 0.931 | 0.803 | 0.973 | 93.35% |
| NN | 0.821 | 0.425 | 0.809 | 0.821 | 0.811 | 0.445 | 0.843 | 82.13% |

**Figure 1.**
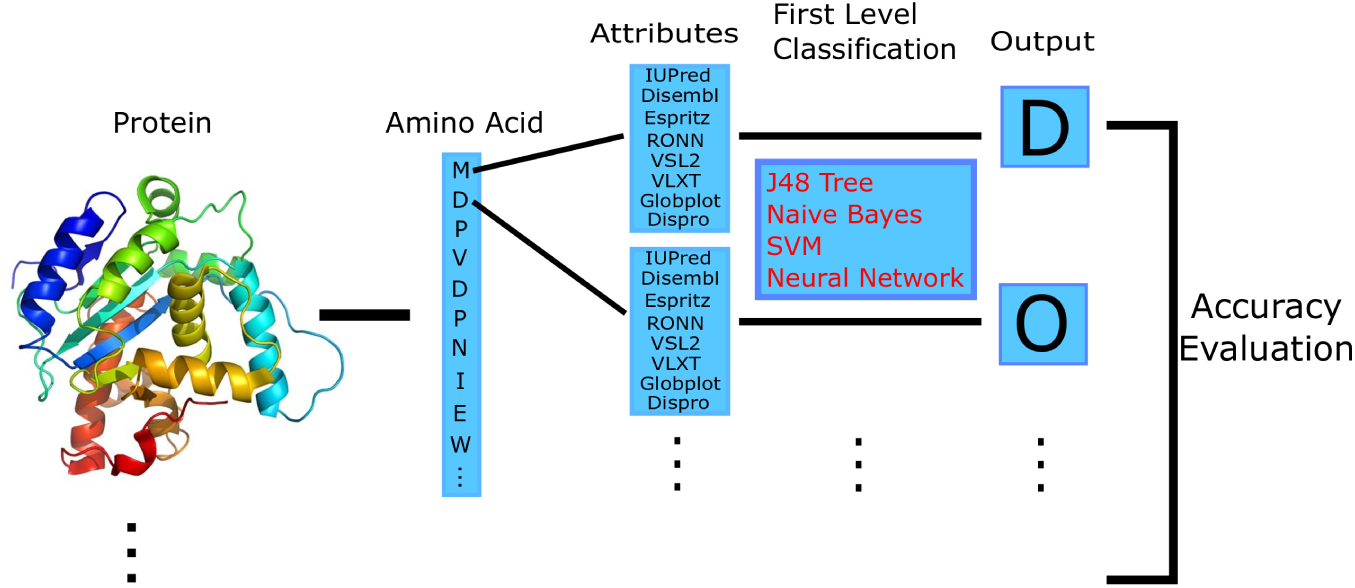
Workflow for Single Amino Acid Approach

**Figure 2.**
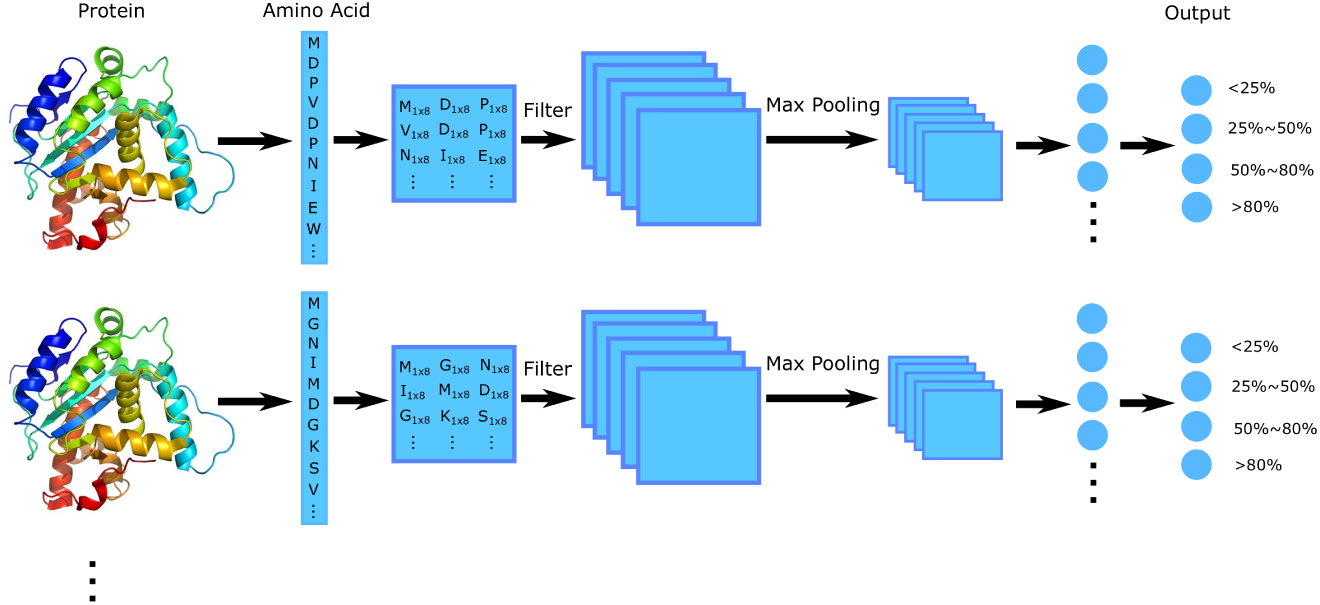
Workflow for Convolutional Neural Network Approach

**Figure 3.**
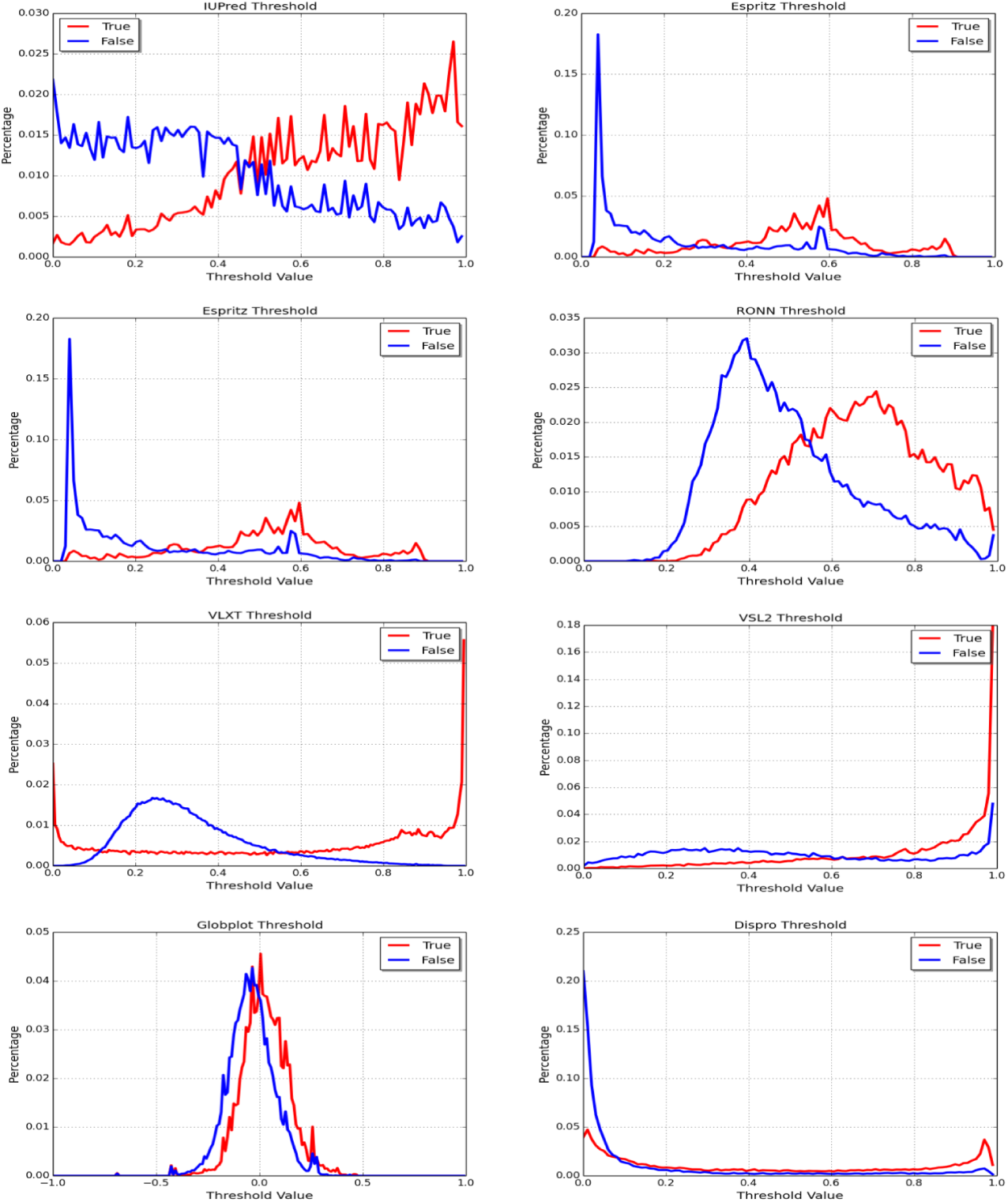
The performance of individual predictors and meta-predictors we used

### The performances of single amino acid classifier

First, We test 4 classifiers,J48, Naive Bayes, Random Forest, Multilayer Perceptron, in Weka with 10 fold cross-validation for all the data we have. The confusion matrix table is shown in Table 2. As this table shows, the correct predict is the number with same results label and ground truth label. The accuracy we get from J48 is 87.6631%. The accuracy of Naive Bayes is 78.0744%. For Random Forest model, the accuracy we get is 93.3497%. The last single amino acid classifier we test is Multilayer Perceptron, the accuracy we get is 82.1334%.

To evaluate these four classifiers together, we compare the accuracy of each classifier while we change the data size. Figure 4 show the accuracy of all four classifiers. We can see that as the size of training data set increases, the accuracy also increases. Among the four classifiers, Random Forest has the highest accuracy, then is J48 Decision Tree, then is Multilayer Perceptron(NN), and Naive Bayes has the lowest accuracy. The reason Naive Bayers is the worst performance among these four classifiers could be that the Naive Bayes is good for handling independent attributes, whereas in our dataset, the attribute are somehow correlated.

**Figure 4.**
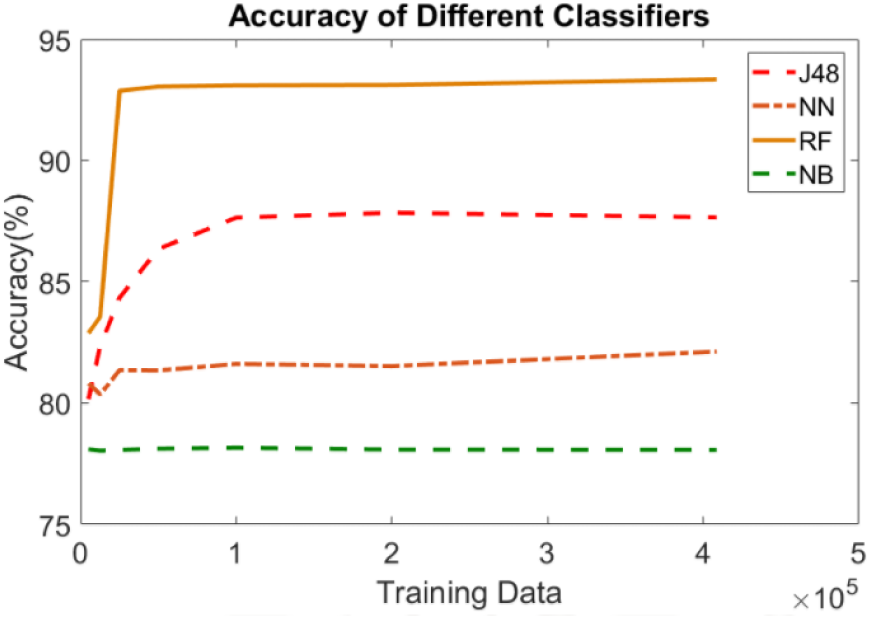
Accuracy Comparison Among Four Classifiers.

Aside from just compare the accuracy, we also use other metrics to compare them, such as false positive rate, F-Measure, MCC and ROC area. The comparison result is shown in Table 3. The FP rate and TP rate are common measures in machine learning, the TP rate is the higher the better and the FP rate is the lower the better. Precision is defined as the number of true positives(TP) over the number of true positives(TP) plus the number of false positives (FP). Recall is defined as the number of true positives (TP) over the number of true positives (TP) plus the number of false negatives (FN). The F-Measure is a measure of a test’s accuracy and is defined as the weighted harmonic mean of the precision and recall of the test. The Matthews correlation coefficient (MCC) is used in machine learning as a measure of the quality of binary (two-class) classifications, the value are more approximate to positive 1 represents a perfect prediction. The ROC curve is created by plotting the true positive rate against the false positive rate at various threshold settings. We can see that Random Forest has the highest value of MCC, ROC, F-Measure, etc. This demonstrates that Random Forest has the best performance among all four classifiers.

### The Performances of whole protein CNN classifier

To build the model, we use both DisProt and single chain PDB training data set for training. 3967 protein sequences are used in CNN model training. The test data is the PDB X-ray dataset as we mentioned in section 2. There are 10525 amino acid chains for testing. Each protein is converted to a matrix of amino acids as shown in Figure 2. In addition, each amino acid is further represented by the eight attributes we used in the single amino acid classifier. All the proteins are classified to four classes as we mentioned in section 2. The result is evaluated as the total correct classification instances among all cases in the test data. After training, we choose the PDB dataset We constructed and test a series of different CNN architectures by changing three parameters: the number of filters, the number of layers and the unit number of fully connected hidden layers. The results are shown in Table 4.

**Table 4.** The accuracy and training time of different CNN models

| Convolution Layers | Number of Filters | Fully connected units | Training Time | Accuracy |
| --- | --- | --- | --- | --- |
| 1 | 8 | 512 | 1112.018s | 93.252% |
| 1 | 16 | 512 | 1331.798s | 95.012% |
| 1 | 8 | 1024 | 1465.740 | 94.127% |
| 2 | 8,16 | 512 | 997.634s | 95.620% |
| 2 | 8,16 | 1024 | 1321.231s | 92.201% |
| 2 | 2,4 | 512 | 563.512 | 86.708% |
| 3 | 8,16,32 | 512 | 523.51 | 72.808% |

We can see that we got highest accuracy of 95.62% when there is 2 convolutional layers with 8 and 16 filters, 512 units in hidden layer, and 0.5 dropout rate at first hidden layer [21]. The training time is 997.634 seconds for building the model on GAIVI cluster. We find that when we increase the convolutional layers the training time is not increased and may decrease. 1 convolutional layer with 8 filters needs 1112.018 seconds to build the model but 2 layers with 8 and 16 filters need only 997.634 seconds. This is because we use 2×2 max pooling layer to reduce the matrix size after each convolutional layer. So with 1 convolutional layer, the matrix will be at 60×50. But with 2 convolutional layers, the matrix size will be 30×25. Since the data size decrease, the training time will not increase when we add convolutional layers.

## Discussion

Currently, there are more than 70 predictors created to predict the IDP. However, all of them have the limitation on N- and C- terminus prediction. In addition to the differences of outputs from predictors, the outputs include both single amino acid classifier and fraction of disorder amino acid in whole proteins. Different outputs are used in different research directions based on whether the goal is to study the biological systems and protein functions. Our study uses a meta-strategy to combine the current predictors and improve the prediction accuracy. Specifically, the CNN model shows highest prediction for IDPs. Since CNN considers the nearby input element by using a filter to affect the output after convolutional layer, this implies a combination of consideration of nearby amino acids could improve the accuracy. The present study is limited in that it requires the quality of data that is collected by the existing prediction tools. As we Table 3 shows, these predictors has accuracy ranges from 64.5% to 76.8%. Since we use eight attributes as the total data for our whole protein disorder prediction, the CNN model learn the pattern of these attributes and automatics amplify or reduce the significant of some attributes. It is important for us to know under each condition we should trust which predictor’s score. Thus, the future work will focus on understanding why the CNN model has an accuracy of 95.62% by combining all these eight predictors.

## Conclusion

In this project, we first proposed single amino acid approach. In this approach, we treat each amino acid as an instance and use four different classifiers, J48, Naive Bayes, Random Forest, and Multilayer Perceptron, to run in Weka with 10 fold cross-validation. We evaluated which classifier has the best performance for single amino acid classification. From our results, we conclude that Random forest has the best performance in single amino acid classification, which has accuracy of 93.35%. Then we proposed Convolutional Neural Network. We choose four labels for each protein to do whole protein classification. When using 2 convolutional layers with 8, 16 filters each layer and 1 fully connected layer with 0.5 dropout, we get the accuracy of 95.62%, which is the best result we get. Our study shows a machine learning based meta-strategy can improve the prediction of intrinsically disorder protein.

## Methods

### Datasets preparation

In this study, we use three independent datasets from DisProt (http://www.disprot.org) and RCSB Protein Data Bank (PDB) database [22].

The first training set is composed from the Database of Protein Disorder (DisProt) (http://www.disprot.org), version 7 v0.3 of September 26, 2016 [23]. The sequences were clustered by CD-HIT with 30 percent sequence identity [24]. The reason we use 30% identity as cutoff value is because usually researchers treat sequences less than 30% identity as functional and evolutionary unrelated. 802 protein sequences with true disordered amino acid residues compared with all residues 92360/408691 were obtained.

The second training set is obtained from RCSB Protein Data Bank (PDB) database [22] of February 28, 2017 by choosing single chain and more than 15 models structures. Structures having ligands or disulfide bonds were removed from dataset. The same sequences with training set were removed, then remaining sequences were clustered by CD-HIT with 30% sequences identity. By applying the distribution of RMSD value of each residue, threshold value 5 was selected as cutoff to determine whether this residue is disordered or structured. From the calculation, 3165 chains have been obtained and the number of true disordered residues compared with all residues in this dataset is 37189/327443. The independent testing set is comprised by PDB X-ray dataset, which was updated on March 2, 2017. After removing the same sequences with both training set and first testing set, the remaining dataset contains 10525 chains and 2579528 amino acid residues. We treated missing residues in PDB file as experimental validated disorder residues because it is hard to obtain the structure of disorder residues. The number of true disordered amino acids in this dataset is 143860.

To preprocess the data, each amino acid is predicted by the available predictors. Every predicted value is consider as an independent variable for that amino acid instance. Each amino acid is labeled as “D” or “O” according to the DisProt and PDB database. “D” means this amino acid is disordered or in disordered region. “O” mean this amino acid is ordered and in well structured region.

### Logistic regression model to select individual predictors

As we mentioned above, we start from a pool of 72 disorder protein predictors that can be searched in PubMed and Web of Science. The first screen step of individual predictors is through literature review and web server checking. We set two criteria for these protein: 1) The predictor is widely used and cited; We choose only predictors that has original paper cited more than 10 times per year. 2) The predictor has well maintained User-friendly web server. From all 72 predictors, we selection 11 predictors as shown in Table 1 for logistic regression model step by step selection.

In order to select the best predictor to include in machine learning model. We test combination of different logistic models using step-by-step forward parameter selection. We use PROC LOGSELECT to test model shown in equation 1 in SAS 9.4 (University of South Florida Academic License.) In this model, *P*(*D*) is the probability of a single amino acid classified as disordered amino acid. P(O) is 1 − *P*(*D*), which is the probability of a single amino acide classified as ordered one. *β*_0_ to *β*_11_ are model estimations. *x*_1_ to *x*_11_ are predict scores for each amino acid instances. In order to get most significant model, we choose the more convergent dataset DisProt that has 408691 amino acid instances as we mentioned in section 2.1.

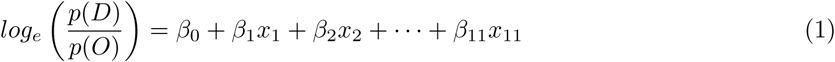

### Single amino acid classifier

As we mentioned in section 3.1, each amino acid of protein are labeled by “D” and “O”. Therefore, single amino acid approaches were first proposed using Weka with 10-fold cross-validation. Here, each amino acid was treated as an instance. There are 408,691 instances in training set, containing 8 attributes. In our single amino acid approach, we use 4 different classifiers in Weka with 10 fold cross validation. The process of our single amino acid approach is shown in Figure 1.

In the single amino acid approach, we use 4 different classifiers, which are Decision Tree-J48, Naive Bayes, Random Forest and Multilayer Perceptron. Here we will just give a brief description of each classifiers.

We first use Decision tree-J48, it is the implementation of algorithm ID3 (Iterative Dichotomiser 3) developed by the WEKA project team. J48 is a popular algorithm used to generate a decision tree developed by Ross Quinlan. Decision trees require relatively little effort from users for data preparation. In order to classify a new item, it first creates a decision tree based on the attributes from the training dataset and predicts the value of a target variable.

The Naive Bayes classifier works well for independent attributes, which is based on the Bayes rule of conditional probability. It will consider each the attributes separately when classifying a new instance.

Random Forest is an ensemble machine learning method for classification, which constructs multiple decision trees at training stage. It is fast and accurate and is better compared with J48 on a single decision tree. Random forest also corrects the overfitting problem of decision tree. It can used some trick to perform better, such as, using bagging to create training set each time and randomly pick some features and then use information gain to pick the best.

Multilayer Perceptron is a type of neural network that usually consists of at least three hidden layers. The node in each layer uses nonlinear activation function. It is different from linear perceptron, in that between the input and the output layer, there can be one or more non-linear layers, called hidden layers. It uses back-propagation in the training process.

### Protein convolutional neural network

To classify the whole protein, we choose 4 labels for each protein based on the ratio of discorded region according to the standard of classification of IDP protein [25]. As shown in Figure 2, if the protein has more than 80% disorder regions, it is labeled as class 1, highly disorder protein. And 50% to 80% disorder regions is labeled as class 2, medium disorder protein. 25% to 50% is labeled as class 3, low disorder protein. Less than 25% is labeled as class 4, ordered protein.

We use a 2-D CNN to take the input of the protein data. For each protein, we extract the eight attributes of every amino acid and fit them into a matrix. Because the length of the protein ranges from 50 to more than 2000. We choose the vector size as 12000 which is enough to contain 1500 amino acids’ data. If the protein is shorter than 1500, we use padding to make the data vector at 12000. If the protein is longer than 1500, only first 1500 amino acids are included into the data vector. After the data vector is constructed, we reshape the data to a 2-D 120×100 matrix for CNN input.

The process for whole protein CNN is shown in Fig 2. In CNN, we first use several filters to extract feature maps of the original data. Then we use max pooling layer to reduce the data dimension. After one or several CNN/max pooling layers the data go through classify layer which is fully connected neural network. The final class can be read from the classifier output. During our experiments, we test different architectures of CNN by tuning the convolutional layers, filters, hidden layers. We test accuracy and training time for each model. All the CNN models are built with Tensorflow flame work on University of South Florida GAIVI GPU cluster with GTX TITAN X as our training hardware.

## Ethics approval and consent to participate

Not applicable.

## Consent for publication

Not applicable.

## Availability of data and materials

The DisProt Dataset can be downloaded from DisProt download page. (https://www.disprot.org/download). The RCSB Protein Data Bank(PDB) dataset can be found from PDB database (https://www.rcsb.org/downloads). The CNN model is built in tensorflow frame work (https://www.tensorflow.org/). The source code is published on author’s github (https://github.com/brandonfire/ProteinDisorder Tensorflow).

## Author’s contributions

LS provided the initial idea and the resources. CY and JJ collected the data and performed the data preprocessing. TZ conducted the initial Weka machine learning experiments. CH and YQ wrote the source code for the machine learning experiments and collected the results. LS, ZH, and CH wrote the paper. All authors contributed to the data interpretation and approved the final manuscript.

## Acknowledgements

Not applicable.

